# The gene annotated by the locus tag At3g08860 encodes a β-alanine/L-alanine aminotransferase in *Arabidopsis thaliana*

**DOI:** 10.1101/576041

**Authors:** Anutthaman Parthasarathy, Francisco C. Savka, André O. Hudson

## Abstract

The aminotransferase gene family in the model plant *Arabidopsis thaliana* consists of 44 genes some of which remain uncharacterized. This study elucidates the function of an uninvestigated aminotransferase annotated by the locus tag At3g08860. The cDNA was shown to functionally complement two *E. coli* mutants auxotrophic for the amino acids β-alanine (non-proteogenic) and L-alanine (proteogenic). The elucidation of At3g08860 activity has the potential to facilitate experiments for the optimization of plant lines involved in nitrogen utilization efficiency, response to hypoxia, osmo-protection, vitamin B5 and coenzyme A metabolism.

## INTRODUCTION

Aminotransferases or transaminases (EC 2.6.1.X) are pyridoxal-5’-phosphate (PLP) dependent enzymes that catalyze reversible reactions between amino acids and alpha keto acids by transferring an amine group from a donor to an acceptor. These enzymes function via a bi-molecular double displacement ping-pong mechanism where an amino acid usually serves as the amino donor and a α-keto acid serves as the amino acceptor (Nelson and Cox, 2000). Aminotransferases are ubiquitous in the three kingdoms and life and are involved in a variety of metabolic pathways including amino acid metabolism, nitrogen assimilation, gluconeogenesis, and responses to a number of biotic/abiotic stresses, among other pathways (de Sousa and Sodek, 2003; Liepman and Olsen, 2004; Rocha et al., 2010; McAllister et al., 2013). The genome of the model plant *Arabidopsis thaliana* contains 44 annotated genes as part of the aminotransferase gene family. In this family, 8 of the 44 genes are annotated as putative alanine aminotransferases. The loci tags are At2g13360, At4g39660, At2g38400, At1g23310, At1g70580, At1g17290, At1g72330 and At3g08860 (Liepman and Olsen, 2004; Niessen et al., 2012). In plants, the biosynthesis of non-proteogenic amino acid β-alanine can be anabolized from four different precursors: (1) the polyamines spermine and spermidine, (2) the nucleotide base uracil, (3) propionate and (4) L-aspartate. However, only the propionate pathway involves a β-alanine aminotransferase (Fig 1A). In contrast, L-alanine is synthesized by the transamination of pyruvate, where L-glutamate serve as the amino donor. This reaction is catalyzed by the enzyme alanine aminotransferase (Fig 1B).

**Fig 1.**
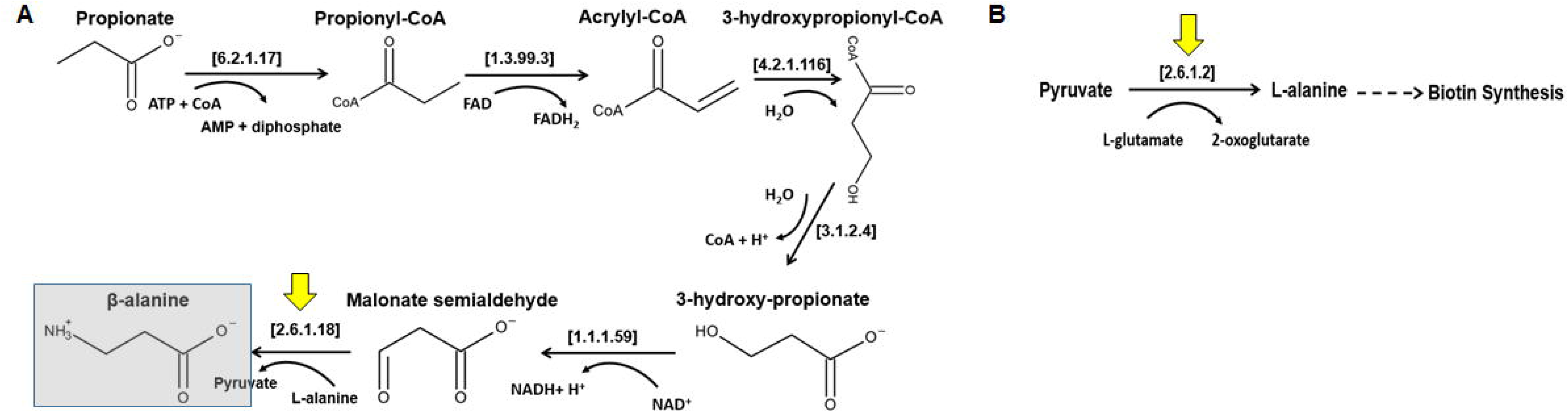
(A) β-alanine synthesis from the propionate pathway. β-Alanine aminotransferase catalyzes the synthesis of β-alanine and pyruvate using L-alanine as the amino donor and malonate semialdehyde as the amino acceptor. (B) Pyruvate is transaminated by alanine aminotransferase to synthesize L-alanine using L-glutamate as the amino donor. The EC numbers of the enzymes shown in brackets correspond to the following enzymes: 6.2.1.17 = propionyl-CoA synthetase, 1.3.99.3 = acyl-CoA dehydrogenase, 4.2.1.116 = 3-hydroxypropionyl-CoA dehydratase, 3.2.1.4 = 3-hydroxyisobutyryl-CoA hydrolase, 1.1.1.59 = 3-hydroxypropionate dehydrogenase, 2.6.1.18 = β-alanine-pyruvate aminotransferase and 2.6.1.2 = alanine aminotransferase. The yellow arrows indicate the reactions catalyzed by β-alanine and L-alanine aminotransferase.

In plants, alanine aminotransferases are important because they are involved in a number of important pathways. For example, it was identified in *A. thaliana* and *Oryza sativa* that two mitochondrial L-alanine/glyoxylate aminotransferases link glycoxylate oxidation to glycine formation (Niessen et al., 2012). The phenotypes of various alanine aminotransferase overexpressed in the *A. thaliana* Col-0 background and in the alanine aminotransferase (At1g17290 and At1g72330) knockouts suggests that nitrogen use efficiency (NUE) could be improved in plants by the overexpression of alanine aminotransferases (McAllister and Good, 2015). Gene regulation studies of alanine aminotransferase in response to low-oxygen stress, light and nitrogen has been studied in many plants and it was shown that hypoxia induced the expression of two distinct alanine aminotransferase genes (At1g17290 and At1g72330) in *A. thaliana* (Miyashita et al., 2007). The function of the gene product from the locus tag At3g08860 has not been experimentally elucidated. Here we present data to show that the gene product of one of the putative alanine aminotransferase genes annotated by the locus tag At3g08860 encodes a β-alanine/L-alanine aminotransferase using an *in vivo* functional complementation experiment.

## Materials and Methods

### Plant Growth and Conditions

*A. thaliana* Col 7 from the Arabidopsis Biological Resource Center (ABRC) was grown on Murashige and Skoog (MS) medium with a 16-hour light (light intensity was approximately 120□Em^−2^□s^−1^) and an 8-hour dark period, with temperatures of 24°C during the light period and 20°mC during the dark period.

### RNA Isolation from Arabidopsis thaliana

Total RNA was isolated from 7 day old Col 7 *A. thaliana* seedlings grown on MS medium using TriZol reagent (Life Technologies). One hundred milligrams of seedlings was ground in liquid nitrogen and homogenized in 1□mL of TriZol, followed by incubation at room temperature for two minutes. Total RNA was precipitated using 1□mL 100% (v/v) isopropanol. The RNA pellet was washed three times with 1□mL 75% (v/v) ethanol. The air-dried RNA pellet was resuspended in 30□μL of Diethyl Pyro carbonate (DEPC)-treated water and quantified using a NanoDrop spectrophotometer.

### cDNA Synthesis

The Reverse Transcription System Kit (Promega) was used to synthesize a cDNA library following the manufacturer’s protocol. One microgram of total RNA from 7 day old seedlings was used to synthesize cDNA. The reaction contained 1μL oligo-dT primer, 1□μg total RNA, 1μ□L of 10_mM dNTP mix, and DEPC-treated water up to 13□μL. The mixture was incubated at 65°C for 5 minutes followed by an incubation on ice for 5 minutes.

### Amplification and Cloning of At3g08860 cDNA

The protein full length protein encoded by At3g08860 is predicted to be 481 amino acids in length. The protein was predicted to be localized to the mitochondria using the TargetP and SUBA subcellular localization prediction tools (Emanuelsson et al., 2007; Heazlewood et al., 2007). The first 93 nucleotides of the full length cDNA was predicted to encode the signal sequence that denote localization to the mitochondria. As such, the first 93 nucleotides was excluded when cloning the cDNA. The At3g08860 cDNA was amplified via PCR. The PCR (50 μL) reaction contained 1 μL (12 pm/ μL) each of the forward primer 5’-CACCATGTCCTCCGTCCGCGAGACCGAGACCGAA-3’ and the reverse primer 5’-*CTGCAG*TCACATCTTGGACATGGCGTGATCCATCAC-3’, 1 mM MgSO4, 0.4 mM of each of the four deoxynucleotide triphosphates, 2 μL of cDNA library and 1 unit of platinum *Pfx* DNA polymerase (Invitrogen Corporation, Carlsbad, CA, USA). The following PCR conditions were used: 1 cycle at 94°C for 3 minutes, followed by 35 cycles of 94°C for 30 seconds, 60°C for 30 seconds, and 72°C for 2 minutes and an indefinite soak of 4°C. The cDNA amplicon was ligated into the pET100/D-TOPO vector (Invitrogen Corporation, Carlsbad, CA, USA). The fidelity of the of the pET100D::At3g08860 construct was confirmed via Sanger nucleotide sequencing using T7 promoter (5’-TAATACGACTCACTATAGGG-3’) and the T7 terminator (5’-TATGCTAGTTATTGCTCAG-3’) primers located on the pET100/D-TOPO plasmid backbone.

### Plasmid for Functional Complementation

The plasmid pBAD33::At3g08860 used for functional complementation experiments of the *E. coli* mutants auxotrophic for β-alanine *(panD)* and L-alanine *(HYE032)* was constructed by subcloning Xba1, Pst1 sites from the pET100D::At3g08860 construct into pBAD33 (Guzman et al., 1995).

### Functional Complementation

Auxotrophic *E. coli* mutants for L-alanine synthesis *(HYE032)* (avtA::GM, yfbQ::KM, yfdZ::FRT, Ala^−^) was obtained from Dr. Dr. Hiroshi Yoneyama from Tohoku University (Yoneyama et al., 2011). β-alanine synthesis *(panD)* (F-, *Δ (araD-araB)567, ΔpanD748::kan, ΔlacZ4787c* (::rrnB-3), *λ^−^, rph-1, Δ (rhaD-rhaB)568, hsdR514)* was obtained from the Coli Genetic Stock Center (CGSC #8404) (http://cgsc2.biology.yale.edu/) (Baba et al., 2006). The auxotrophic strains were transformed with pBAD33 or pBAD33::At3g08860. Transformants were selected on LB media supplemented with chloramphenicol (34 μg/mL). Colonies were then replica-plated on M9 agar plates containing M9 salts (1X), 2 mM MgSO_4_, 0.1 mM CaCl_2_, and 0.1% glycerol (w/v), +/− 0.2% glucose or arabinose, +/− β-alanine/L-alanine (10 μg/μL). In testing of the *panD* (β-alanine auxotroph), uracil was also required (10 μg/μL).

## RESULTS AND DISCUSSION

Literature mostly supports the idea of alanine accumulation during hypoxia (unknown reasons) and an increase in alanine aminotransferase activity as plants return to normoxia (de Sousa and Sodek, 2003). This is perhaps a mechanism for maintenance via an increase of the nitrogen pool/skeletons, since the assimilation of inorganic nitrogen affects anaerobic tolerance (Miyashita et al., 2007). During hypoxia/anoxia in plant tissues, fermentative products such as acetaldehyde, ethanol, and lactate can accumulate where the regeneration of NAD^+ by^ lactate dehydrogenase and alcohol dehydrogenase enhances seedling survival (Ismond et al., 2003). In fact, in *A. thaliana*, it was reported that pyruvate decarboxylase was specifically induced during oxygen limitation, but not other stresses (Kürsteiner et al., 2003). An alternative way to counter hypoxia would be through alanine aminotransferase, which could reduce the flux of carbon through lactate (which is acidic and has the potential to regulate the cytoplasmic pH) and prevent the buildup of toxic acetaldehyde (Ricoult et al., 2006). It appears that alanine fermentation primarily functions to regulate the level of pyruvate. Pyruvate is not only a known activator of the alternative oxidase (Vanlerberghe et al., 1999), but has also recently been shown to interfere with the hypoxia-induced inhibition of respiration (Gupta et al., 2009; Zabalza et al., 2009). Therefore, in order to control the rate of respiratory oxygen consumption when the oxygen availability is low, it is important to prevent pyruvate accumulation. Alanine fermentation has the potential to accomplish pyruvate accumulation with the additional advantage that alanine can accumulate to high concentrations without the detrimental side effects that go along with the lactate or ethanol fermentation pathways (Rocha et al., 2010). Alanine enters the propionate pathway of β-alanine synthesis via the enzyme β-alanine aminotransferase [EC 2.6.1.18], exchanging an amino group with malonate semialdehyde, and generating pyruvate and β-alanine (Fig 1A).

Functional complementation assays showed that the plasmid harboring the *At3g08860* cDNA was able to rescue both the L-alanine and β-alanine auxotrophs. The assay show that both *E. coli* mutants were able to grow on β-alanine and L-alanine free media compared to the vector only control, which needed both amino acids to grow (Fig 2A and 2B). Interestingly, in the *panD* background, the cDNA rescued growth under repressible conditions (plus glucose) and not under inducible conditions (plus arabinose) (Fig 2A) whereas the opposite is true in the *HYE032* background (Fig 2B). This result suggests that the enzyme is probably involved in β-alanine metabolism and not L-alanine metabolism. The results of the assays demonstrated a definitive preference of enzymatic activity towards the synthesis of β-alanine suggesting that At3g08860 is maybe involved in osmo-protection, formation of vitamin B5 and coenzyme A, given the fact that β-alanine is involved in these pathways.

**Fig 2.**
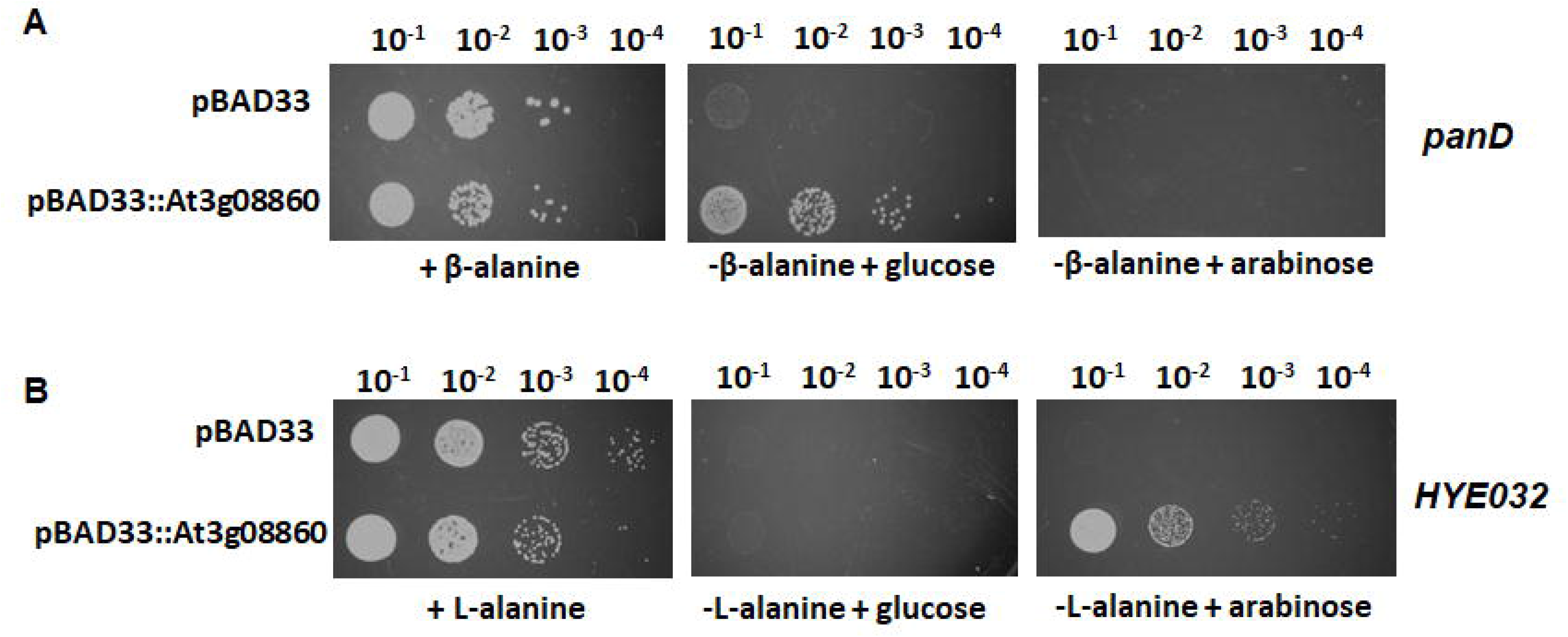
Functional complementation assay. (A) Functional complementation of the *panD E. coli* mutant, which is auxotrophic for β-alanine. (B) Functional complementation of the *E. coli HYE032* mutant, which is auxotrophic for L-alanine. The plasmids used were pBAD33 and pBAD33::At3g08860. Transformants harboring pBAD33 or pBAD33::At3g08860 were grown to an OD of 1.0 measured at 600nm. The strains were serially diluted to 10^−1^, 10^−2^, 10^−3^, and 10^−4^ using 0.85% (w/v) saline. Five□μL was replica plated on M9 medium with or without β-alanine or L-alanine supplemented with 0.2% (w/v) arabinose or glucose.

Studies investigating genes (across a wide array of metabolic/cellular processes) have identified the *At3g08860* locus as responsive to changes in light, which indirectly could affect carbon/oxygen availability/concentration (Thum et al., 2008). Previous work suggested that the protein is localized in either the mitochondria or the peroxisome (Niessen et al, 2012). The visualization of cellular localization using GFP-tagged transcripts was unsuccessful, however based on its sequence; this aminotransferase is believed to be localized in the mitochondria. This suggests an involvement in photorespiration, particularly as it relates to glycine synthesis (following glycolate oxidation to form glyoxylate) as detailed by Niessen et al., 2007, who also demonstrated that alanine:glyoxylate aminotransferase activity was the only aminotransferase activity detected within the mitochondria (Niessen et al., 2012). They also demonstrated that alanine degradation resulted in an increase in CO_2_ release following addition of alanine to mitochondrial extract, implying that alanine degradation increased photorespiratory activity (Niessen et al., 2007). Further suggesting the involvement of At3g08860 in photorespiration based on the gene expression in hydathode tissue (Wenzel et al., 2008).

## ACKNOWLEDGEMENTS

We thank Dr. Hiroshi Yoneyama from Tohoku University for providing the *E. coli HYE032* mutant. We would like to acknowledged the College of Science and the Thomas H. Gosnell School of Life Sciences at the Rochester Institute of Technology for ongoing support.

